# Production and Characterization of Monoclonal antibodies to Xenopus proteins

**DOI:** 10.1101/2022.12.06.519341

**Authors:** Brett Horr, Ryan Kurtz, Ankit Pandey, Benjamin G Hoffstrom, Elizabeth Schock, Carole LaBonne, Dominique Alfandari

**Affiliations:** The University of Massachusetts Amherst, Dept of Veterinary and Animal Sciences; Antibody Technology Resource, Fred Hutchinson Cancer Research Center, Seattle, WA; Northwestern University

**Keywords:** Xenopus, Monoclonal Antibody, Hybridoma, Method

## Abstract

Monoclonal antibodies are powerful and versatile tools that enable the study of proteins in diverse contexts. They are often utilized to assist with identifying subcellular localization and characterizing the function of target proteins of interest. However, because there can be considerable sequence diversity between orthologous proteins in Xenopus and mammals, antibodies produced against mouse or human proteins often do not recognize Xenopus counterparts. To address this issue, we refined existing protocols to produce mouse monoclonal antibodies directed against Xenopus proteins of interest. Here we describe several approaches for the generation of useful mouse anti-Xenopus antibodies to multiple Xenopus proteins and their validation in various experimental approaches. These novel antibodies are now available to the research community through the <u>D</u>evelopmental <u>S</u>tudy <u>H</u>ybridoma <u>B</u>ank (DSHB).

**Summary statement:** The manuscript describes the generation and characterization of novel monoclonal antibodies to *Xenopus laevis* proteins using refined hybridoma production methods suitable for basic science research labs.

## Introduction

Monoclonal antibodies (mAb) are produced by fusing B-lymphocytes to immortal myeloma cells (Kohler and Milstein, 1975). Successful fusion creates a hybridoma cell line that produces a single heavy and light chain antibody specific to a particular antigenic determinant (epitope) of a given antigen. Like its myeloma parent, this cell line is immortal and can be propagated indefinitely. It can also be frozen, subsequently thawed, and expanded to produce large amounts of monospecific antibodies. Here, we make use of this powerful and versatile technology focused on monoclonal antibodies generated for the study of *Xenopus laevis*, a vertebrate model system widely used for cell and developmental biology research.

The Xenopus model system has been at the forefront of cell and developmental biology for decades and has led to the discovery of critical developmental pathways that define the induction of embryonic tissues, the formation of primary axes, and the regulation of the cell cycle (Heasman, 2006). The large size of Xenopus embryos (~1mm) and the relative ease with which hundreds of synchronously developing embryos can be obtained make Xenopus ideally suited for studying the function and composition of protein complexes during early development (Exner and Willsey, 2021; Kostiuk and Khokha, 2021; Medina-Cuadra and Monsoro-Burq, 2021; Niehrs, 2022). Advances in genome annotation and genomic methods have allowed quantification of mRNA expression at most stages of embryo development and in most cell types (Gilchrist et al., 2020; Lindeboom et al., 2019). Advances in proteomics have also provided similar insights into protein expression and post-translational modifications at many developmental stages (Lindeboom et al., 2019; Lombard-Banek et al., 2019; Saha-Shah et al., 2019; Wasson et al., 2019). While these discovery approaches are powerful, building upon them with mechanistic studies requires antibodies specific to individual proteins that can be used in studies to determine subcellular localization, confirm knockdown efficiency, and identify protein complexes formed during specific developmental stages.

While some proteins, including histones and many kinases, are highly conserved between Xenopus and human, the average conservation of protein sequences between human and Xenopus is much lower (68%), which results in poor commercial monoclonal antibody cross-reactivity. Some approaches have taken advantage of the amenability of Xenopus embryos to large-scale protein purification to generate antibodies against proteins expressed in specific tissues or subcellular compartments (Nakazato and Ikenishi, 1989; Sakakibara et al., 2005). For example, hybridomas libraries were produced against proteins isolated from both Xenopus and Pleurodeles oocyte nuclei and screened for their ability to recognize specific structures associated with Lampbrush-chromosomes (Lacroix et al., 1985; Roth and Gall, 1987).

While polyclonal antibodies are relatively easy and inexpensive to make, they are a finite resource that should be carefully characterized due to variability in specificity between each batch. Examples abound in which a new lot of a commercial antibody no longer recognizes the original, specific target protein or begins to recognize additional, nonspecific targets. Consequently, recent efforts focus on clonal antibody production via either hybridoma fusion or using methods where immunoglobulin genes are cloned and expressed in cell lines (Ouisse et al., 2017).

Here we describe a monoclonal antibody production pipeline utilizing “classical” fusion methods paired with refinement strategies and recent advances in cell biology and cell culture. We outline both low throughput techniques applicable to most laboratories as well as automated robotic improvements for medium-scale studies.

## Results and Discussion

Given the growing number of monoclonal antibodies and the convention of naming them according to the plate number and position (e.g., 9E10 for myc), we chose nomenclature that includes initials before the plate number, position and the target protein name (e.g., DA5H6sox3). All antibodies generated in this study are distributed via the Developmental Study Hybridoma Bank (DSHB https://dshb.biology.uiowa.edu) at cost.

### Antigen production

We have used three strategies to produce antigens: bacterial fusion proteins, proteins produced and purified from Human Hek293T cells, and peptides. While we were able to reliably produce mAb with both bacterial fusion proteins and Hek293T proteins, we did not have any success with the peptides. For our initial antibody production, each protein antigen was a full-length protein tagged with FLAG. This was justified, as we could rapidly obtain these constructs from individual investigators in plasmids such as pCS2 that allow both mRNA production via SP6 RNA polymerase and expression in cells by transfection via the CMV promoter. Following transfection in Hek293T cells, a typical construct would yield, on average, ten micrograms of protein per microgram of DNA. Thus, transfecting between five and ten 10 cm plates yielded sufficient protein for a full immunization schedule (200 to 1000μg of protein). Xenopus XTC cells were also transfected with the same constructs which enabled us to screen hybridomas by indirect immunofluorescence. This initial approach worked well for many proteins (**Table 1**), and typically, four mice were injected with five proteins each, using either Freund or Adjuplex adjuvants. While certain antigens were dominant in multiply immunized animals, reactivity against all five proteins was often found in the serum (Fig.1). In general, the mice immunized with Freund’s adjuvant gave a much stronger immune response than those immunized with Adjuplex, but both adjuvants generated useful hybridomas. To conserve limited resources, often only the “best mouse” (defined by strong and broadly reactive serum titers to the target proteins) was selected for fusion rather than doing side-by-side comparisons. We observed that in many cases, the variability of responses between mice was more significant than the variability between adjuvants, highlighting the difficulty of pinpointing “ideal” protein/adjuvant/animal combinations. Nevertheless, each immunization produced hybridomas to nearly all targets that could recognize the protein by ELISA, Western blot, and immunofluorescence when tested on cells overexpressing the specific target. In addition, we also isolated two anti-FLAG hybridomas that are useful to the community.

**Table 1.**
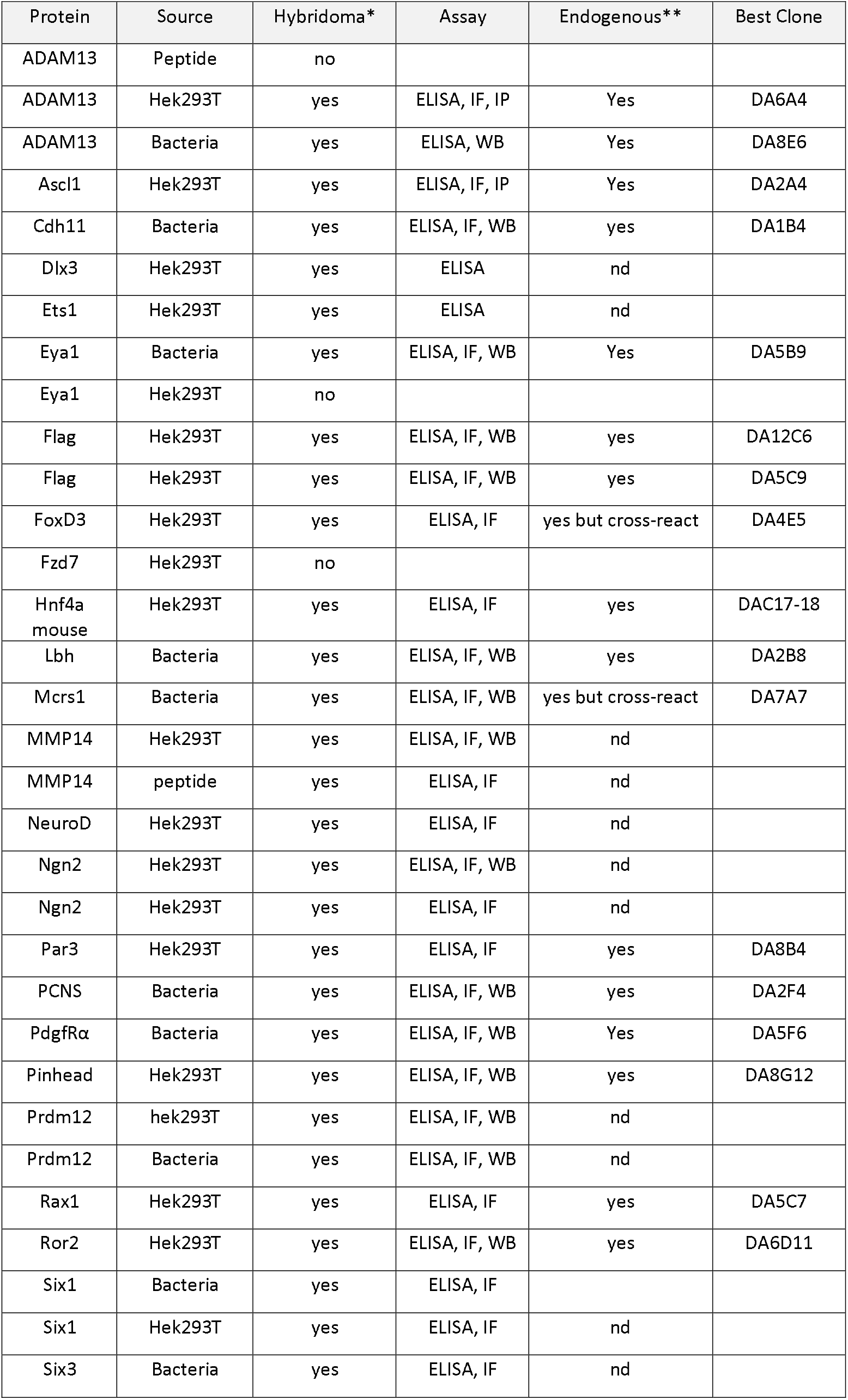
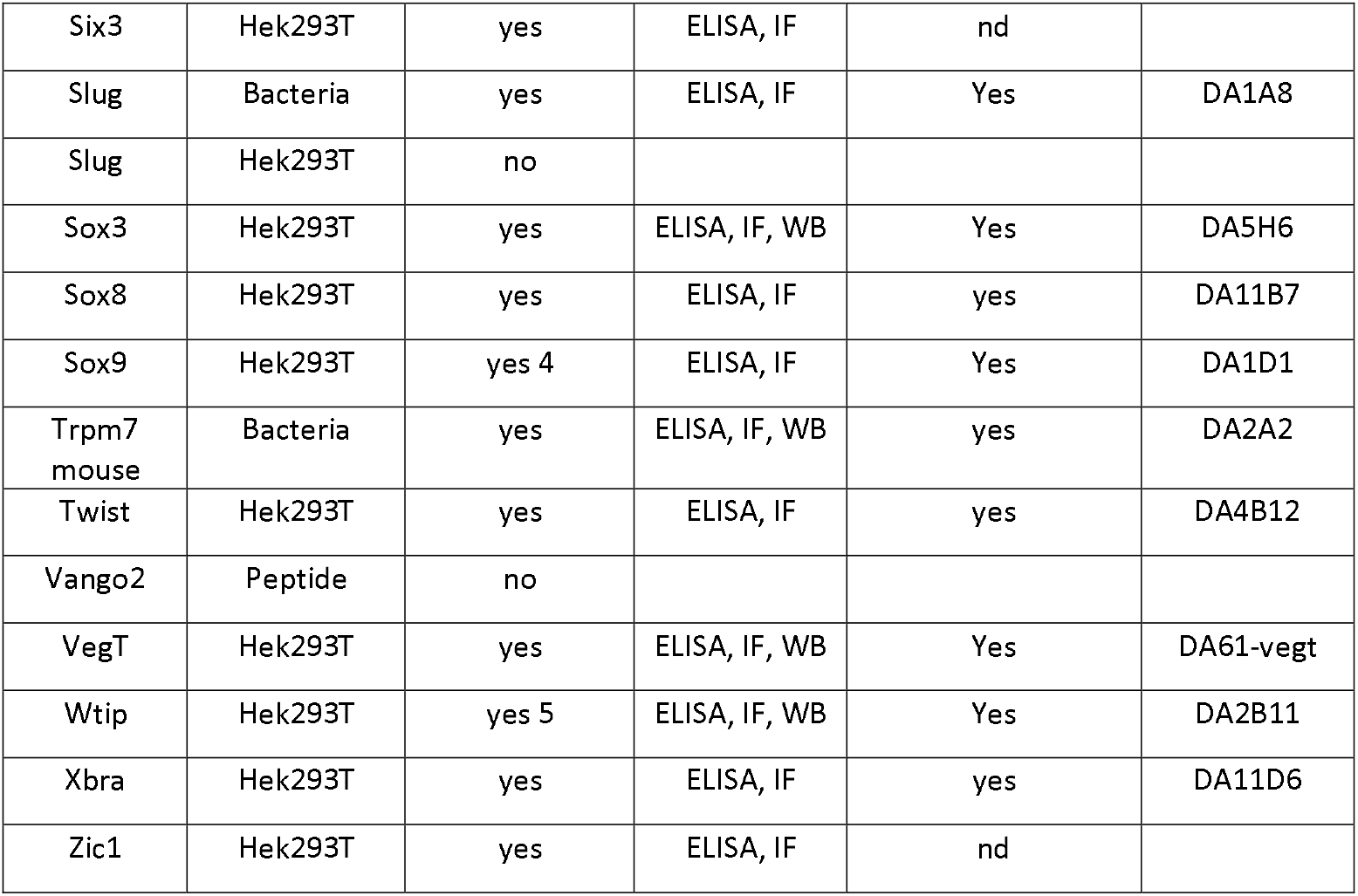
Table of antigens used to generate hybridomas. The protein name and source of the antigen (Hek293T, Bacterial fusion protein or peptide) are provided. *The table indicates whether a stable hybridoma was identified and subcloned (Yes/No). Assays in which the hybridomas were tested (ELISA, Immunofluorescence IF, Western Blot WB). ** Indicates if the antibody recognizes the endogenous target in Xenopus embryos (Yes or not determined nd). Best clones indicate that the hybridoma was selected, subcloned, frozen, and distributed.

**Figure 1.**
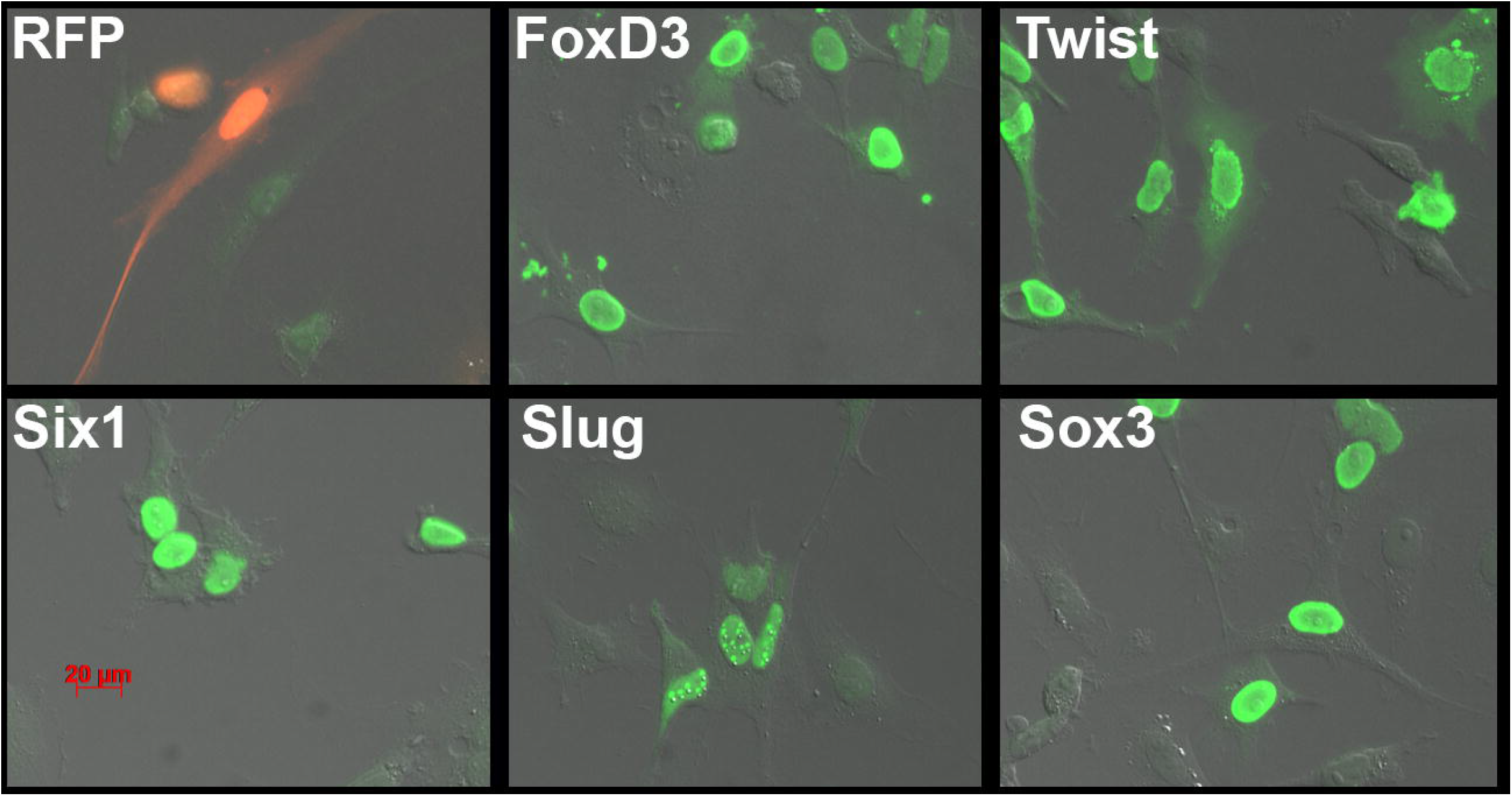
Immunofluorescence testing of serum. The serum from mouse M3 immunized with Foxd3-Flag, Twist-Flag, Six1-Flag, Slug-Flag, and Sox3-Flag produced in Hek293T cells was tested by immunofluorescence on XTC cells transfected with each of the target or RFP-Flag as a negative control. A secondary Alexa488-anti-mouse antibody was used to visualize the mouse primary antibodies (Green). Notice the green nuclei in each of the transfections, with some non-transfected cells lacking signal.

### Limitation of the Hek293T production approach

While we were successful in generating many useful hybridomas using full-length proteins produced in Hek293T cells, there were cases where this proved problematic. First, we found that some proteins could not be produced or purified in sufficient quantity to be used for immunization (e.g., Adam11). Second, we found that for some proteins that are members of multiprotein families with significant sequence homology (e.g., FoxD3), all clones exhibited broad cross-reactivity with other Fox transcription factors. To circumvent these issues, histidine-tag fusion proteins were generated using less conserved regions of these factors. These proteins were then expressed in and purified from bacteria. This approach was utilized for several transcription factors, including Slug, Twist, Six1, Prdm12, Eya1, Six3, and Mcrs1. This approach was also used to purify the cytoplasmic domains of transmembrane proteins (PCNS, PDGFRα, ADAM13). For the most part, these proteins generated strong immune reactions and yielded many hybridomas. On the other hand, bacterial fusion proteins often carried with them contaminants that were highly immunogenic and could hijack the reaction of the immune system. To avoid the detection of hybridomas that produce antibodies toward bacterial contaminants co-purified with the fusion proteins, we used extracts from Hek293T transfected with the full-length proteins for the ELISA assays (See Primary Screening).

We found that, independently of the method of production and purification, we were able to produce hybridoma that specifically recognized the purified antigen. The complete list of antigens produced, the source of production, and the identification of positive hybridomas is provided in **Table 1**.

### Other antigens

We also tested three other types of antigens that will not be described in detail here. First, we used peptides corresponding to optimal sequences based on antigenicity and exposure on the folded protein coupled to Keyhole limpet haemocyanin (KLH). Unfortunately, while we did get strong immune reactions from the injected mice, we did not identify useful hybridomas. We also produced mRNA and immunized mice using lipid nanoparticles as previously described for the coronavirus (Gebre et al., 2022). In effect, *in vitro* transcribed mRNA is assembled into modified lipid vesicles that are used to deliver the mRNA intramuscular so that the foreign protein is produced *in situ*. As noted previously, most laboratories working with Xenopus embryos routinely inject *in vitro*-transcribed mRNA, making this approach extremely attractive. Unfortunately, the immunization of four mice with this strategy did not generate any immune response to the targets (Xenopus Adam9, Adam11, and Adam19). Finally, we also used Xenopus embryo extracts and Xenopus cell lines for immunizations. These gave solid immune responses and multiple hybridomas, but the targets were challenging to identify using mass spectrometry.

### Hybridomas production

While our initial fusions were performed using the classical PEG fusion with success, we moved to electrofusion, given its significantly higher efficiency (Twenty-fold). In practice, this means that one mouse with an average spleen containing 100 million splenocytes that yielded an average of 1000 hybridomas with PEG instead yielded 20,000 hybridomas with electrofusion. While screening 20,000 hybridomas for five targets is not routinely practical in a small lab, this allowed us to aliquot and freeze splenocytes for fusion at a later time should good hybridomas not be identified in the initial screening. It also meant that if one fusion was contaminated, we could easily repeat that fusion using frozen splenocytes and not lose the weeks of immunization. In general, two fusions were performed for each set of antigens using 10 million splenocytes and 10 million Sp2/0 per fusion that was then split into four 96-well plates. On average, each well contained 1 to 3 clones (400 to 1200 hybridomas/fusion).

### Primary Screening

To further optimize the pipeline, supernatants from 4 or 8 plates (one or two fusions) were pooled for the primary screen to facilitate rapid identification of positive hybridomas with a minimum number of ELISA and immunofluorescence plates. In practice, at day ten post-fusion, 30μl was collected from each well and pooled into a “master plate”, preserving the well identity (All A1 wells would be A1 in the master plate see diagram Fig.2). While this can be done manually, we utilized an Opentrons robot (OT2, https://opentrons.com) to increase efficiency and decrease human error. Depending on the number of hybridomas per well, the supernatants were pooled from either four plates (if more than three clones per well) or eight plates (if fewer than three clones per well). The master plate was then used to apply the supernatant to ELISA plates coated with individual proteins as well as glass bottom plates containing XTC cells transfected with plasmids coding each antigen (Fig.2) paired with fluorescent markers. We assigned fluorescent markers as follows: our first construct was transfected with nuclear cherry, second with membrane cherry, third with nuclear sirius, fourth with cytoplasmic sirius, and fifth with RFP. To facilitate screening, we typically paired nuclear fluorescent markers with transmembrane proteins and membrane fluorescent markers for nuclear proteins so that signals could readily be detected with a fluorescent microscope set on a triple filter channel (Zeiss Axiovert 200M, Red/Blue/Green). Once transfected individually, cells were pooled and plated in glass bottom 96-well plates. Immunofluorescence was carried out using an Alexa-488 anti-mouse secondary antibody. Once positive wells were identified, for example, A6, that position was then tested on each fusion plate (1A6, 2A6, 3A6, 4A6) to identify the exact position of the positive colony (deconvolution, Plate/Line/Column). In most cases, one well produced supernatant that recognized the same protein by both ELISA and immunofluorescence. In some cases, we identified wells that recognized all the proteins in both assays. These turned out to be directed against the FLAG-epitope tag used in the purification. We selected and characterized two of these anti-FLAG antibodies that are now freely available (DSHB https://dshb.biology.uiowa.edu/search?keywords=flag).

**Figure 2.**
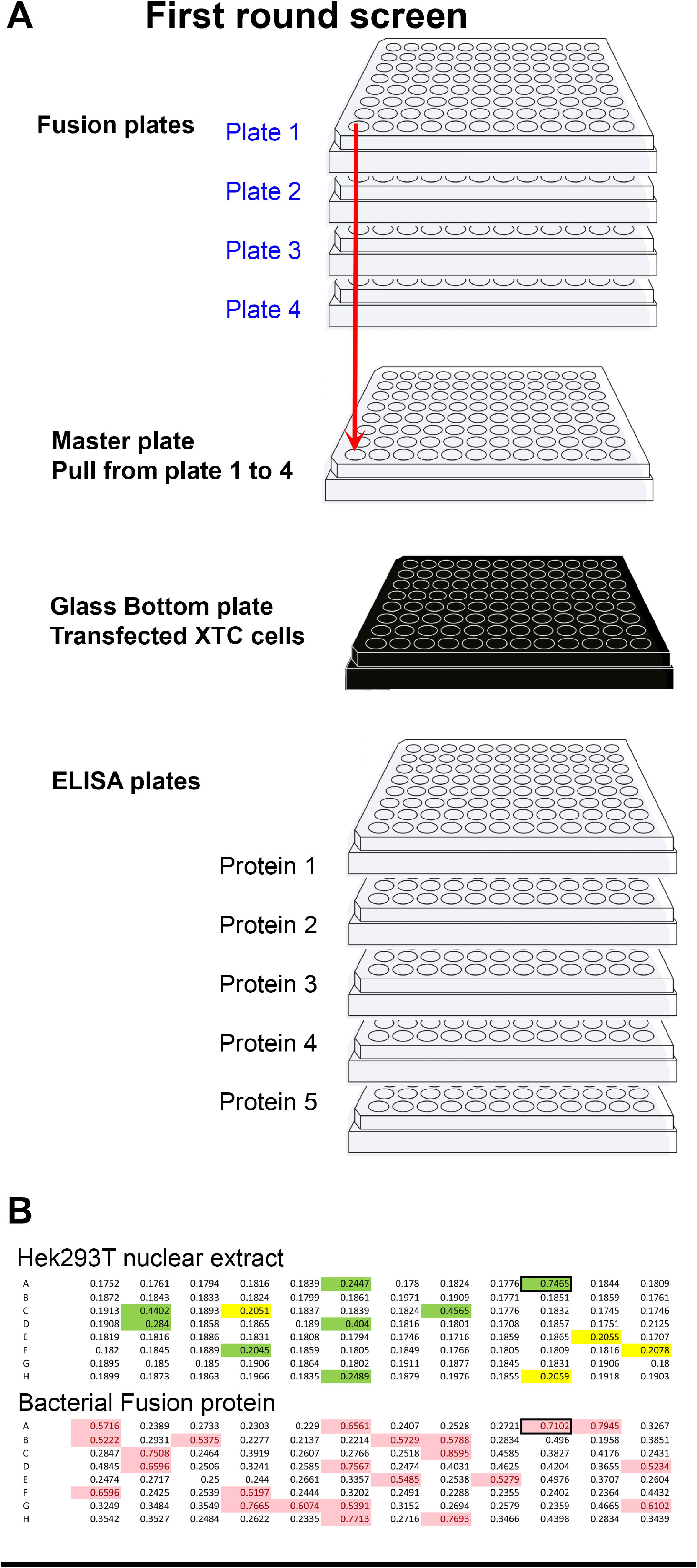
Primary screening method. A) Diagram representing the strategy used to rapidly identify wells producing antibodies to multiple targets at once. From top to bottom, four plates per fusion were used. Thirty microliters of each well were harvested and placed in the master plate (total volume 120 μl). The master plate is diluted with 200 μl of TBST. Fifty microliter was then distributed into one glass bottom plate containing XTC cells transfected with each of the five targets. Simultaneously, 50 μl of supernatant is added to each well of five ELISA plates, each coated with individual proteins. Wells positive in either or both assays (A1 to H12) are then tested on each of the original plates (blue) to identify the plate number (deconvolution, 1A1 to 4H12). B) primary screen by ELISA using the cytoplasmic domain of Adam13/33 expressed in Hek293T cells (upper) and his-tag bacterial fusion protein (lower). The mice were immunized using the bacterial fusion protein. Highlighted wells are above the chosen background (0.2 for Hek293T and 0.5 for bacterial fusion protein). Wells highlighted in green are common to both ELISA and were selected for further screening.

When a western blotting antibody was preferred by an investigator, the primary screen was performed using ELISA, followed by a Western blot on the protein expressed in Hek293T cells instead of immunofluorescence. This method prioritized the identification of antibodies to linear epitopes of denatured proteins rather than native conformations in fixed cells.

For the hybridomas that were produced against bacterial fusion proteins, we screened using the same proteins expressed in Hek293T cells. This was critical when screening master plates, where every well corresponds to a pool of 4 wells, each with three hybridomas (12 potential colonies). In such cases, 75% of the colonies that appeared to detect the bacterial fusion protein did not detect the protein when expressed in Hek293T cells (Fig.2 B). This suggested that the hybridoma either recognized one of the plasmid expression-tags (S-tag or his-tag in pet30a) or a co-purified bacterial protein contaminant. Being able to eliminate 75% of false positive clones was essential for the successful identification of useful antibodies (in this example, 8 hybridomas). For ELISAs, we found that protein purification from Hek293T cells was often not necessary. Instead, in this case, we either used raw nuclear extract (NE-per) or total protein extract (in PBS) to coat ELISA plates. This was especially convenient for proteins that were poorly expressed and/or purified. In the absence of detergent, the transfected proteins were sufficiently enriched to produce a detectable signal above the background.

### Clone isolation

Once a positive well was identified, it was a race against time to identify the colony that produced a useful antibody. A typical well had three colonies of various sizes growing. In general, 50% of hybridomas do not produce immunoglobulin due to the random distribution of chromosome sets during fusion. These tend to grow faster as they are not using their resources for antibody production. As cells multiply, the border between colonies becomes less obvious, and if refeeding is necessary, cell mixing becomes a real possibility. To maximize efficiency, we performed several rounds of screening from the original well to confirm the usefulness of the hybridoma before cloning (see secondary screening below). We then hand-picked individual colonies from each positive well (Fig.3). This step was critical, as it minimized the chances of losing the clone that was producing the antibodies of interest. In practice, a typical first-round screening identified 10 to 50 hybridomas per fusion. At three colonies per well, this led to 30 to 150 new wells after picking, a manageable number for the next series of screening. In the alternative scenario, if ultimate dilution was used directly, it would lead to 10 to 50 new 96-well plates, a number that far exceeds the capacity of most small labs. Another advantage of manually picking colonies is that the transferred colony will produce new supernatant ready to use within three to five days with minimum contamination from the original well (1/200μl) that still contained the antibody produced during the first ten days. Furthermore, by transferring the larger hybridomas to new wells, this technique provided a chance for smaller hybridoma colonies to grow to a size that allowed picking a few days later if none of the previously picked colonies produced the antibody of interest.

**Figure 3.**
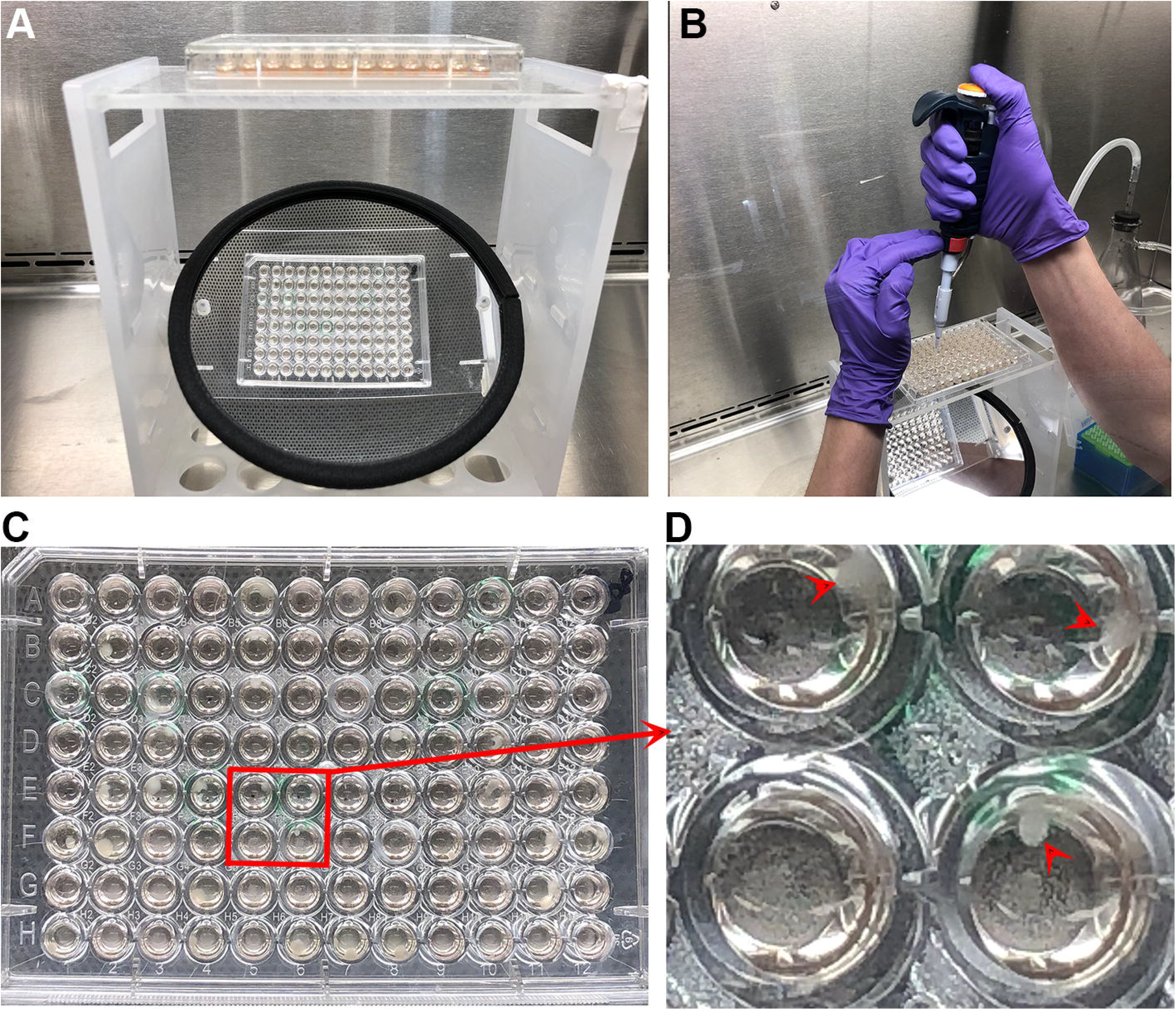
Manual clone picking. A) Plates containing the hybridoma fusion were placed on a mirror to visualize large colonies. B) Investigator pipetting a single colony guided by the mirror. C) Photographs of the plate as seen in the mirror. D) Higher magnification of 4 wells. Individual colonies are highlighted with a red arrowhead.

Antibodies recognizing the full-length proteins expressed in cell lines by immunofluorescence and/or western blot were frozen and shared with individual investigators for further testing using the endogenous proteins and in the context of knockdown. In general, while we were able to produce monoclonal antibodies that recognized each of the targets that were used in our immunization (148), only a fraction (approximately 17%) were sufficiently selective or sensitive to detect the endogenous proteins (25). Antibodies that do not recognize the endogenous protein can still be used in transfection experiments or through overexpression to perform western blot, immunofluorescence, and co-immunoprecipitation without the requirement for tags that may interfere with the function of the protein.

### Secondary screening; Confirmation of Specificity in vivo

As our main goal was to produce antibodies that could recognize endogenous proteins expressed in Xenopus embryos, each positive hybridoma required further characterization. Promising antibodies were screened for their ability to detect endogenous protein by western blot, and in some cases, they were further characterized by immunostaining or immunoprecipitation. Examples of each assay are given below.

### Western blotting

We utilized a capillary western blotting system (WES ProteinSimple^™^) in which 24 individual hybridoma supernatants can be tested at once using only 10μl of supernatant per sample. This meant that hybridomas could be tested before subcloning using the small amount of supernatant from the original well identified after deconvolution. Hybridomas were screened for those that could detect the over-expressed protein in Hek293T cells and a similar size band in embryo lysates prepared from a stage at which the mRNA was expressed while detecting no band at stages in which the mRNA was absent (https://www.xenbase.org/entry/). In cases where a morpholino was available to deplete the protein of interest, we tested protein extracts from non-injected control embryos compared to embryos injected with ten nanogram of the relevant morpholino. In our experience, this dose was sufficient to effectively block translation in all but the most abundant proteins (e.g., lbh)(Cousin et al., 2011; Weir et al., 2021). An example of this strategy is presented in Figure 4 for the Xbra (Tbxt) monoclonal antibody (Fig.4A-C).

**Figure 4.**
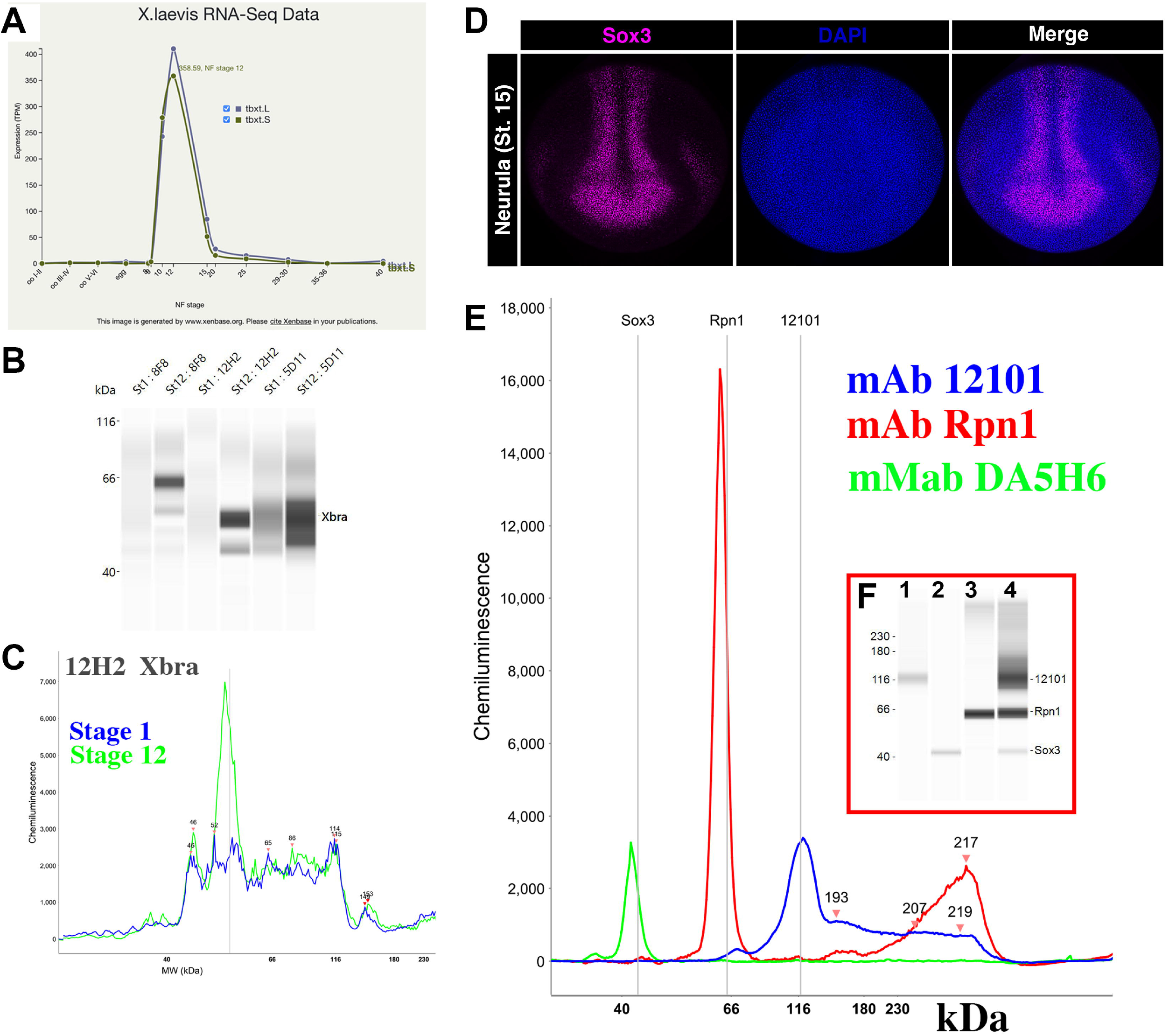
Secondary screening. A) RNAseq expression data for Xenopus *brachyury (tbx?)* from Xenbase. There is no mRNA for this gene at stage 1 (fertilized egg), while it is strongly expressed during gastrulation (stage 12). B) Capillary western blot (0.04 embryo equivalent) at stage 1 and stage 12 for 3 independent hybridomas (8F8, 12H2, 5D11). Notice that bands are visible only at stage 12. C) Histogram of the signal intensity of the chemiluminescence for clone 12H2 at stages 1 and 12. Notice that a single peak is present in stage 12 (green) but not stage 1 (blue), while the background is common. D) Whole mount immunofluorescence of neurula stage embryo. Sox3 is in magenta, and DAPI to stain the nuclei is in blue. MAb DA5H6sox3 detects the nuclei of cells within the neural plate. E) Capillary western blot of embryo extracts (0.04 embryo equivalent per lane) incubated with antibodies to Sox3 (green), Ribophorin1 (Rpn1, red), and the muscle marker 12101 (blue). F) The red insert represents the line view of the chemiluminescence signal obtained (left). Antibodies were incubated either separately (lanes 1-3) or together (lane 4).

This step was critical to identify antibodies that recognized linear epitopes that are masked in the folded protein or in multi protein complexes and might also be masked in immunoprecipitation and immunofluorescence assays.

### Immunostaining

Our original intent was to test all positive clones by whole mount immunostaining. However, we found that both alkaline phosphatase and peroxidase staining required significant optimization for each antibody, making it impractical to carry this out for each clone. Instead, the suitability of antibodies for use in immunostaining assays was tested on a case-by-case basis. Figure 4 shows one such example for mAb DA5H6sox3 directed against the transcription factor Sox3 using immunofluorescence. This antibody readily detects nuclear Sox3 in the neural plate of Xenopus embryos (Fig. 4D). Importantly, it does not cross-react with the related SoxB1 factor Sox2 (not shown), making this a valuable antibody for studies of neural development. MAb DA5H6sox3 also detects endogenous Sox3 in embryo extract (Fig.4E). Similarly, we were able to obtain wholemount immunostaining (Fig.5A) for brachyury (Xbra/Tbxt) and Adam13/33 (Fig.5B). We also found that explants of cranial neural crest cells were an excellent tool to detect endogenous proteins present in these cells (Fig.5C, DA1A8slug).

**Figure 5.**
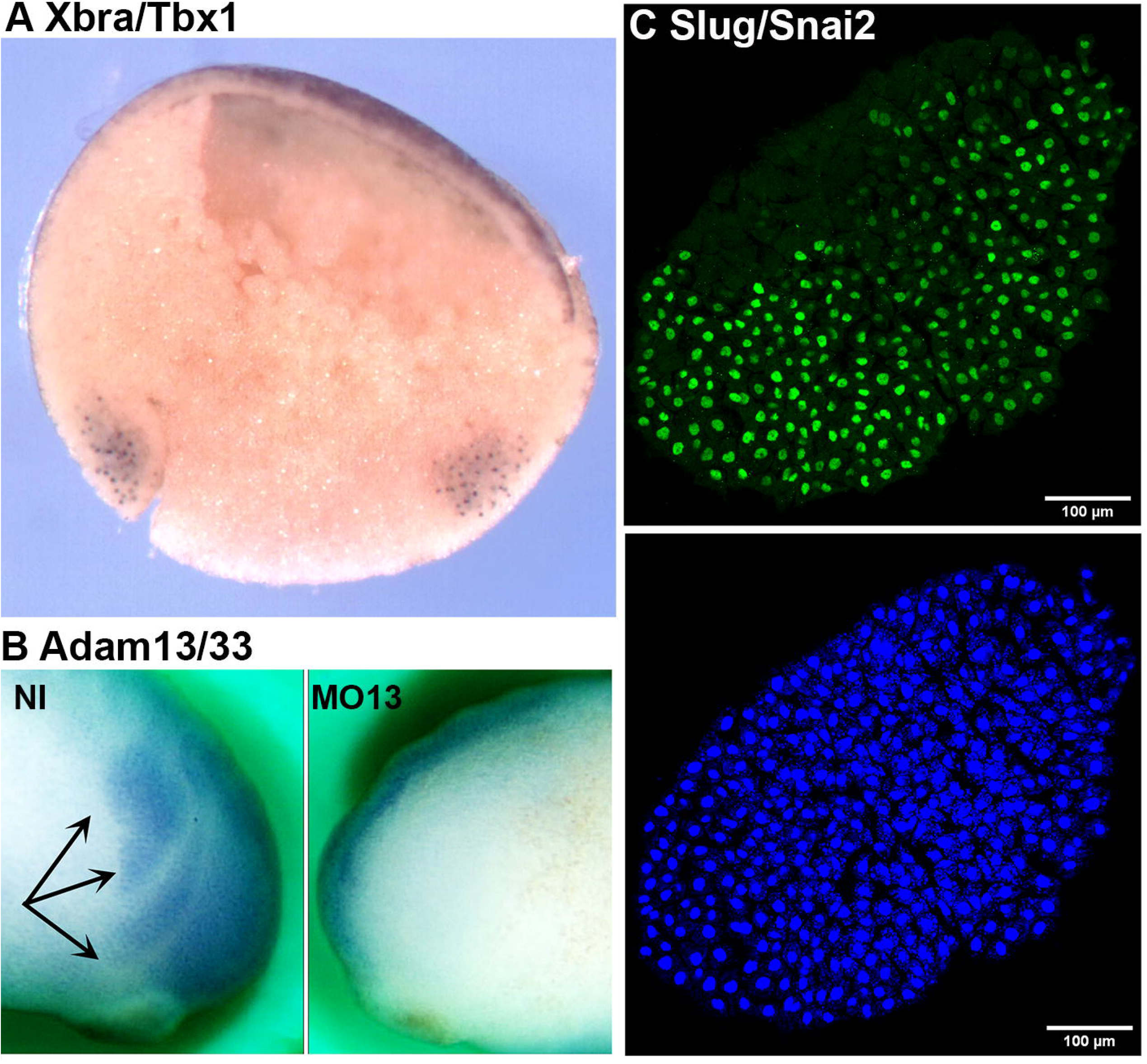
Immunostaining. A) Whole mount immunostaining using DA12H2xbra on bisected gastrula. The dorsal side is to the left, and the animal pole is up. The antibody stains the nuclei from both the dorsal and ventral marginal zones (mesoderm). B) Whole mount immunostaining using a monoclonal antibody to ADAM13 (DA8E6adam13) on embryos injected at the 2-cell stage with a morpholino that blocks translation of Adam13. The cranial neural crest cells (black arrows) are visible on the non-injected side but are absent on the injected side (MO13). C) Immunofluorescence on cranial neural crest explant using mAb DA1A8slug (green) and counter stain with DAPI (blue). Note that most of the nuclei are stained for slug except for a small section on the left of the explant (dorsal) that is likely to be composed of neural plate cells.

### Immunoprecipitation

In cases where a commercial antibody (polyclonal) that recognized the Xenopus protein by Western blot was available, we were able to test the hybridoma of interest for its utility in immunoprecipitation experiments. For these assays, the monoclonal antibody was used to pull down the target protein from an embryo extract which was then detected by western blot using the commercial antibody. Alternatively, immunoprecipitates were tested by tandem mass spectrometry to determine if the expected protein target was pulled down (e.g., mAb Rpn1, data not shown). This step was critical to identify antibodies that may work in chromatin immunoprecipitation (ChIp) assays.

### Conclusion and future directions

Animal model systems are critically important to advance both basic science and biomedical research. The amphibian *Xenopus laevis* has been used for decades, first as a pregnancy test (human HCG induces the laying of eggs) (Elkan, 1938) and then as a model to understand embryo development and evolution. Given the external development (embryos develop autonomously outside of the mother) that allows investigators to view and experiment on each stage, the ability to generate a large number of embryos, and the advance in genomics, Xenopus is now a choice model system to test the effect of known human mutations causing diseases (Moody et al., 2015; Willsey et al., 2022). One of the resources that has lagged is the availability of high-quality monoclonal antibodies to study protein localization and function. Our laboratory has produced more than 100 monoclonal antibodies that recognize 25 endogenous protein targets in the embryo. These antibodies are available worldwide and are adding to the existing list of polyclonal (809) and monoclonal (767) antibodies that have been shown to function in Xenopus (https://www.xenbase.org/reagent). When comparing the *Xenopus tropicalis* proteome (to avoid issues of duplicated genes) to that of humans, we found 36 identical proteins and 865 proteins with over 90% sequence identity (from 11,182 identified orthologues). We selected a cutoff of 90% as it is typically a good indication that epitopes might be sufficiently conserved to provide cross-reacting antibodies. This suggests that the number of cross-reacting commercial antibodies is unlikely to increase in the future unless companies design their epitopes to be conserved across multiple species.

Another approach that has been used extensively in the pharmaceutical industry is the use of recombinant antibodies (Maruthachalam et al., 2022; Panagides et al., 2022). Libraries of variable chain antibodies exist that can be used to screen and identify binders. While this is an extremely efficient way of producing large amounts of a single antibody that can be sold and tailored to a specific application (for example, inserting the variable region into the human constant region), it is not a directly usable resource that can be provided to a typical size laboratory. It also requires the same amount of antigen to screen the libraries. We tested a phage display library to identify targets that would bind specifically to Xenopus proteins. While we performed four rounds of enrichment, we did not find any individual clones that recognized our protein of interest (data not shown). A similar approach using the single-chain camelid immunoglobulin to produce antibodies to Xenopus protein (Itoh et al., 2019) has only shown moderate success as well. As the technology improves, we expect that this will become a good solution for molecular biology laboratories that are used to screening phage libraries to produce their own recombinant antibodies.

## Material and Methods

### SP20 ATCC CRL-1581, SP2/0-Ag14

SP2/0 myeloma cell line was grown from frozen stock in RPMI media supplemented with 10% FBS, 100U/ml Penicillin, 0.1 mg/ml Streptomycin, 2 mM L-glutamine, and 1 mM sodium pyruvate. Cells were grown for up to one week, maintaining a density of less than 1.10^6^/ml. On the day before the fusion, cells were expanded into as many plates as needed with an expected density of 5.10^5^/ml to 1.10^6^/ml.

### Hek293T ATCC CRL-3216

Hek293T cells were grown in RPMI media supplemented with 10% FBS, 100 U/ml Penicillin, 0.1 mg/ml Streptomycin, 2 mM L-glutamine, and 1 mM sodium pyruvate. Cells were passed the day before transfection to 30% confluence and transfected with PEI (1mg/ml in water) at a ratio of 2 million cells, 10 μl of PEI per μg of plasmid DNA. The DNA was mixed in pure RPMI (200 μl/μg of DNA) containing 10 μl of PEI (1 mg/ml). The solution was mixed using a vortexer and incubated at room temperature for 15 min before being added dropwise to plated cells. Cells or supernatant were collected 36 to 48h after transfection and extracted for either ELISA (see below) or protein purification. Most proteins utilized were FLAG-tagged and were purified on FLAG-affinity gel (Abm). Proteins were eluted using 100 mM glycine pH 2.8 and immediately neutralized using 1M Tris pH 9.5 (50 mM final). Proteins were dialyzed against PBS and quantified using a BCA assay (Pierce). One aliquot was run on an SDS-PAGE gel which was subsequently stained with Coomassie to evaluate protein purity.

### Xenopus XTC cells

Xenopus XTC cells were grown in 67% L15 media supplemented with 10% FBS, 100U/ml Penicillin 0.1 mg/ml Streptomycin, 2 mM L-glutamine, 50 μg/ml Gentamycin and 1 mM sodium pyruvate at room temperature (20°C to 25 °C) without CO_2_. Transfection was performed using Fugene HD (Sigma) according to the manufacturer’s instructions. Typically, one well of a six-well plate (60 to 80% confluence) was transfected with 0.5 to 1 μg of DNA mixed with 1.5 to 3μl of Fugene, yielding 30 to 70% transfection efficiency.

### Bacterial fusion proteins Pet30a

Full cDNA or specific regions were selected for their comparative variability and cloned using Infusion (Takara) into bacterial expression plasmid (Pet30a). The constructs were transformed into BL21 DE3 pLysS (ThermoFisher) *E. coli*, grown to 0.5 OD prior to stimulation with 1 mM IPTG and grown for an additional 3 hours, after which the bacteria were pelleted. If the solubility of a fusion protein was unknown, native purification was performed first. The bacterial pellet was resuspended in sonication buffer containing 2 mM PMSF and sonicated three times for 60 seconds on ice. After the addition of Triton X-100 to 1% final concentration, the bacterial lysate was spun for 20 minutes at 10,000g at 4°C to separate soluble and insoluble fractions. The insoluble pellet was then extracted in 8 M Urea containing 100 mM NaH_2_PO_4_ and 10 mM Tris pH 8.0, mixed on a vortex, and rotated for 30 minutes at room temperature (RT) prior to spinning for 20 minutes at 10,000g at RT. The supernatants were incubated with Ni-NTA agarose resin (HisPur, Sigma-Aldritch) overnight at 4°C (native) or 30 minutes at RT (denatured). Beads were washed in sonication buffer containing 20 mM imidazole (native) or 8 M Urea pH 6.3 (denatured). Proteins were eluted with a step gradient of imidazole (100-400 mM) for the native purification or 8 M Urea at pH 4.5 for the denatured proteins. All proteins were dialyzed against PBS and quantified using a BCA assay (Pierce). The purity of fractions was visualized on SDS-PAGE using Coomassie Brilliant Blue (ThermoFisher) before immunization.

### Immunization

Eight-week-old female BalbC mice (Jackson Laboratory) were immunized with either Freund’s adjuvant (Sigma) or Adjuplex (Empirion LLC). The immunization schedule was the same independent of the adjuvant, with one primary immunization on day 1, followed by one boost on day 21. All immunizations were done intraperitoneally with a maximum of 100 μg of purified protein representing either a single protein or 20 μg of up to 5 different purified proteins. Serum was collected on day 30 and tested by western blot against the protein expressed in Hek293T cells and by immunofluorescence against the protein expressed in Xenopus XTC cells. Mice whose serum displayed robust antigen recognition were selected for fusion. If antigen recognition was limited, another boost was performed three weeks after the previous boost. Prior to splenectomy, mice were rested for one month and immunized with the protein in PBS without adjuvant three days prior to sacrifice.

### Fusion

Spleens were harvested aseptically in a biosafety cabinet. Splenocytes were isolated in RPMI media (Cytiva) and washed. Red blood cells were eliminated with RBC Lysis Buffer (Sigma) according to manufacturer’s instructions. Before fusion, SP20 cells were washed with RPMI and pelleted to remove any remaining serum. Initial fusions were performed using PEG (Roche) using up to 150 million splenocytes and 50 million SP20 using a standard protocol as described in (Cousin et al., 2000). Most of our fusions were subsequently performed using electrofusion (BTX, Harvard apparatus) using 10 million splenocytes and 10 to 30 million SP20 per fusion to obtain a similar number of hybridomas (approximately 1000). BTX settings were: 35 seconds AC alignment, 2 pulse at 10 μs at 1000V, 9 seconds post fusion AC, using the Eppendorf isoosmolar buffer. For each spleen, two-thirds of the splenocytes were aliquoted and frozen in 90%FBS and 10% DMSO for future fusions. Each fusion was resuspended in 25 ml of RPMI media containing (20% FBS, HAT Hypoxanthine-Aminopterin-Thymidine), 100 U/ml penicillin 0.1 mg/ml streptomycin, 2 mM L-glutamine, 1 mM sodium pyruvate and Hybridoma Fusion and Cloning Supplement (Roche) in a deep 10 cm plate (Fisher). 1 to 3 days later, the fusion was collected, spun down at 300g for 5 min, and distributed in four 96-well plates with 200 μl of the same media per well.

### Primary Screening

At day ten post-fusion, hybridomas from 24 randomly selected wells were counted, and from there, the total number of colonies was extrapolated. The supernatant was then tested by ELISA on each antigen. For bacterial fusion protein immunogens, the primary screen was performed using the same protein produced in human Hek293T cells. In most cases, Hek293T cells were transfected with the target Xenopus protein (using PEI). The proteins were expressed for 48h, and cells were washed with PBS and frozen. Protein extraction was done using PBS with 5 mM EDTA and protease phosphatase inhibitor complex (Thermofisher) by passing cells through a 24g needle ten times. Insoluble debris was pelleted for 10 min at 16,000g at 4°C. The supernatant containing the transfected protein was directly coated onto ELISA plates without additional purification. For nuclear proteins, nuclear extractions were performed (Pierce NE-per) diluted and used directly in ELISA. Coating efficiency was tested using an anti-FLAG antibody (Sigma M2).

### Secondary screening

The optimal secondary screen was dependent on the desired endpoint application (e.g., Western blot, immunofluorescence, immunoprecipitation). Here two main screens were utilized. The first screen used Xenopus XTC cells transfected with each target and a fluorescent marker (target 1 plus nuclear cherry, target 2 plus membrane cherry, etc.) After transfection, cells were collected, mixed, and plated in 96-well glass bottom plates (CellVis) previously coated with bovine fibronectin (Sigma) at 5 μg/ml. Twenty-four hours after seeding, cells were fixed using 4% paraformaldehyde in 1X MBS for 30 min at room temperature. Cells were permeabilized in 1X MBS containing 0.5% triton X100 for 5 min at RT, washed with PBS, and blocked using PBS containing 1% BSA for at least one hour at room temperature or overnight at 4°C. Plates were kept for up to one week at 4°C. The second screen was performed using an automatic capillary western blotting system (WES, ProteinSimple^™^) following manufacturer instructions.

### Whole mount immunostaining

Fluorescence: *Xenopus laevis* embryos were fixed in MEMFA, bleached, blocked in PBS 1% BSA and 10% goat serum for 1h at room temperature, and incubated overnight at 4°C with the primary antibody (1:100 mAb DA5H6sox3). Embryos were washed and incubated overnight at 4°C with an anti-mouse secondary antibody conjugated to an Alexa Fluor (1:500) and DAPI (1:5000). Embryos were imaged using a Nikon C2 upright confocal. Peroxidase: Xenopus embryos were treated as above. The primary and secondary antibodies (goat anti-mouse HRP 1/1000 Jackson immunoresearch) were incubated overnight at 4°C and washed six times for 10 min at room temperature with PBS 0.1% tween (PTW). KPL TrueBlue substrate was added to the embryos until the label was visible. Overstained embryos were washed in PTW and re-stained until desired contrast was obtained.

### Subcloning

Individual colonies from positive wells of the seeded 96-well plates were picked manually using a P2 pipetman aspirating 1 μl and transferred into 200 μl of media in a new 96-well plate (Fig.1). Briefly, 96-well plates were placed on a mirror in the biosafety cabinet (Fig.1 A), the positive well was identified, and the tip of a P2 pipetman was slowly lowered to cover a single colony, which was aspirated (Fig.1B). The procedure was repeated on each visible colony in each of the positive wells. Subclones were retested 3 to 5 days after transfer, expanded into six-well plates, and subsequently frozen. The best candidates were immediately sub-cloned by seeding one new 96-well plate at 1 cell per well (ultimate dilution) retested, expanded, and frozen. Only hybridomas that went through the two rounds of sub-cloning were considered clonal.

## Acknowledgment

The author would like to thank all Alfandari lab members for their discussion and suggestions. We would also like to thank Dr. Irini Topalidou and Dr. Amy Burnside for their proof reading of the manuscript. We would like to thank Dr. Inchul Yeo for his help in the characterization of the antibody to VegT. We would also like to thank Dr. Sergei Sokol, Takuya Nakayama, and Helen Willsey for their help in the characterization of multiple monoclonal antibodies. We would like to thank all Xenopus contributors that have sent plasmid to produce proteins. We thank the current and former staff at the Fred Hutch Antibody Technology Resource and the University of Massachusetts Amherst IALS microscopy and mass spectrometry facilities for their help.

## Data availability

All antibodies produced are available from the DSHB if fully characterized or by request to Dr. Alfandari at if partially characterized. Antibody descriptions will be linked to gene pages on Xenbase.

